# Mind-manifesting? Examining naturalistic reports from psychedelics, cannabis, meditation, and dreams from the ALTERD mobile application

**DOI:** 10.64898/2026.07.28.741354

**Authors:** Conor H. Murray, Michael Angyus, Sam Suchin, Max Suchin, Martin Monti

## Abstract

Altered states of consciousness (ASC) refer to states that differ markedly from normal waking consciousness. ASC may occur spontaneously, such as in dreaming, or through consciousness modifying techniques, from meditation, to cannabis and psychedelics. The etymology behind the term psychedelic, meaning mind-manifesting, is based on the hypothesis that certain substances, including serotonergic hallucinogens, do not merely distort perception, but also afford genuine observation and insight into the nature of mind, a claim that remains empirically contested today. Here, we address this psychedelic hypothesis using 2,242 reports from the Psychedelics, Cannabis, Meditation, and Dreams categories from the ALTERD mobile application. Across categories, we examined semantic similarity, topic structure, visual phenomenology, and alignment with *a priori* ASC themes. The Psychedelics category was most semantically similar to Dreams, consistent with a hallucinogenic character, yet it also uniquely coupled with Meditation across ASC themes, including Personal Growth, lending support for the psychedelic hypothesis. Cannabis showed the opposite pattern: similar to Meditation semantically, but not aligned on ASC themes such as Personal Growth. Together, these findings indicate that psychedelics induce a dream-like state, yet support a meditation-like trajectory toward personal growth, whereas cannabis may share phenomenological features with meditation, but lacking a transformative therapeutic character.

## Introduction

Altered states of consciousness (ASC) refer to states that differ markedly from normal waking consciousness, occurring spontaneously, such as in dreaming and near-death experiences, or through consciousness modifying techniques, from practices like meditation, to psychoactive substances like cannabis or psychedelics (Dittrich, 1998; Fort et al., 2025; Studerus et al., 2010). Despite the diversity from which altered states of consciousness arise, ASC nonetheless contain shared characteristics that are reproducible across individuals and modifying techniques (Dikovskaya et al., 2025).

ASC are often associated with psychedelics, a term coined by psychiatrist Humphry Osmond from the Greek *psyche* (mind/soul) and *delos* (manifest/reveal). The term was intended to capture a class of substances whose effects “make manifest, open to sight, evident” aspects of nature, including what Huxley termed “Mind at Large” (Huxley, 1954). This etymology encapsulates a theoretical claim – the psychedelic hypothesis – whereby certain substances do not merely distort perception, but also occasion observation and insight into the nature of mind. Critically, this hypothesis stands in opposition to the more conventional view that such substances, while producing perceptual distortions, do not afford any privileged observation or insight, and should be simply referred to as hallucinogens (American Psychiatric Association, 2022).

In recent years, investigators have leveraged online data sources to inform existing nomenclature surrounding psychoactive drugs (Baggott et al., 2010; Martial et al., 2019; Zamberlan et al., 2018) or for testing theoretical frameworks for consciousness and ASC (Fort et al., 2025; Schmidt & Berkemeyer, 2018). However, few sources are currently available to probe phenomenology across varied states of consciousness. ALTERD is a purpose-built mobile application in which users prospectively document first-person experience reports across specific categories, including Psychedelics, Cannabis, Meditation, and Dreams among others. With over 200,000 individual experience reports across 30,000 users, this application has the scale and categorical structure required to map the phenomenology of ASC and to empirically test existing nomenclature and theoretical frameworks. In the current analysis, we used machine learning to examine how the categories relate to one another and to the thematic structure of ASC identified in a recent qualitative analysis (Dikovskaya et al., 2025). To test the psychedelic hypothesis, we used the Dreams and Meditation categories as comparators. Specifically, reports containing similarities to Dreams were interpreted as supporting the hallucinogenic nomenclature, whereas reports containing similarities to Meditation were interpreted as supporting the psychedelic nomenclature. Our rationale for these comparators is that dreaming is inherently illusory, whereas meditation occasions genuine observation and insight related to the mind (Dahl et al., 2015; Dikovskaya et al., 2025; Lutz et al., 2008). Together, our approach aims to test the construct validity of the mind-manifesting label for psychoactive substances by determining whether the phenomenology of ALTERD reports from Psychedelics, or Cannabis, more closely aligns with phenomenology of ALTERD reports from Dreams or Meditation.

## Methods

### Dataset

Data were obtained from the ALTERD mobile application (alterd.com), a purpose-built platform designed to collect first-person reports of altered states of consciousness. From a dropdown menu, users of this platform may self-select into one of nine default experience categories, specifically, “Baseline,” “Tired,” “Exercise,” “Meditation,” “Alcohol,” “Cannabis,” “Psychedelics,” “Dream,” or “Stimulant.” Users also have the option to freely enter text to indicate an additional category of interest, or create a sub-category field, such as “psilocybin.” Following the selection, users are then presented with a note-writing screen to freely enter text associated with their experience. The present analysis focused on four of the default altered experience categories: Dreams (n=831 reports), Meditation (n=2,071 reports), Cannabis (n=18,250 reports), and Psychedelics (n=2,826 reports). For Psychedelics, we determined the number of reports which were associated with a text field sub-category, specifically, psilocybin/mushroom (n=828 reports), LSD/acid (n=300 reports), DMT (n=119 reports), ketamine (n=56 reports), and salvia (n=10 reports). Similarly, for Meditation reports, user entered sub-categories included breath techniques (n=78), music meditation (n=25), Vipassana (n=11), Mindfulness (n=10), Transcendental (n=9), and Loving-kindness (n=7) among others. The categories were also examined in relation to the Baseline category, which is designed to collect user experience prior to reporting altered state experiences (n=18,179 reports). All reports were submitted voluntarily and anonymously. The UCLA IRB determined this work is not research involving human subjects as defined by DHHS and FDA regulations and therefore full IRB review and approval was not required for the present analysis.

### Preprocessing

Report-level analysis: At the level of the full report, the raw text was filtered to a minimum of 100 words to exclude text fragments, and duplicate reports were excluded. In Supplemental methods, we developed a filter to examine the extent to which the 100-word threshold removed “unsuitable” reports for the analysis to validate this approach. Because the Cannabis category was substantially larger than the others after filtering (n=4,006), it was randomly downsampled to the median size of the remaining three categories (random_state=42). In Supplemental methods, we applied the same suitability filter on the resulting Cannabis sample to confirm this sample was not by chance relatively “unsuitable.” Final analytic sample sizes were: Dreams n=361, Meditation n=641, Cannabis n=620, Psychedelics n=620 (total N=2,242 reports). For word cloud visualizations (see Supplementary Material), standard English stopwords and contraction fragments (e.g., ‘s’, ‘t’) were additionally removed, with a minimum word length of three characters applied (Dikovskaya et al., 2025).

Sentence-level analysis: Visual scene sentences were extracted from each report using a keyword filter targeting sensory and spatial language. Extracted sentences were filtered to a minimum of 50 words. Unlike at the report level, Cannabis sentences were not downsampled to probe the full breadth of cannabis-related visual scenes, and the full set of extracted sentences was retained across all categories for sentence-level analyses.

### Text Embedding

All text was converted to vector representations using the all-MiniLM-L6-v2 model (Reimers & Gurevych, 2019), implemented via the SentenceTransformers library in Python 3.11.14. The all-MiniLM-L6-v2 model transforms text into lists of 384 numbers so that texts with similar content are represented by similar vectors in a 384-dimensional space. Embeddings were computed at two levels: at the report level, where each full report was embedded as a single vector, and at the sentence level, where each sentence associated with visual scenes were embedded separately.

### Dimensionality Reduction and Visualization

Report embeddings were projected to three dimensions using Uniform Manifold Approximation and Projection (UMAP; (McInnes et al., 2018)) to visualize the global structure of the consciousness state space. The three emergent axes were phenomenologically labeled by submitting 8 reports from the extreme low and high ends of each axis (48 reports total) to Llama 3 8B (Grattafiori et al., 2024), running locally via Ollama, with the prompt: “Read both sets and propose a concise phenomenological label for this axis in the format: [low pole]

↔ [high pole].” Sentence embeddings were similarly projected to three dimensions using UMAP with identical hyperparameters. To ensure all reports were represented regardless of length, UMAP was fitted on a stratified sample of 3,000 sentences sampled across reports, then applied to the full sentence set.

### Pairwise Category Similarity

At the report level, a 4×4 pairwise cosine similarity matrix was computed across all report embeddings, with all embeddings L2-normalized and scaled to unit length to enable consistent angular comparison prior to computation. This analysis was conducted on both the full between-subjects dataset (N=2,242) and a within-subjects subset consisting of 21 participants who contributed at least one report in each of the four categories (N=84 reports, one report randomly sampled per category per person). At the sentence level, an analogous 4×4 similarity matrix was computed, but due to the relatively high number of sentences, instead of computing all pairwise sentence similarities, we first computed the mean embedding of all sentences within each category to form a single category centroid (L2-normalized), and pairwise cosine similarity was then computed between these four centroids.

### Topic Modeling and Topic Similarity

Per-category topic modeling followed Ermakova et al. (2025) (Ermakova et al., 2022). At both the report and sentence levels, embeddings were reduced to two dimensions using UMAP and clustered using Hierarchical Density-Based Spatial Clustering of Applications with Noise (HDBSCAN; (Campello et al., 2013)). Each topic was characterized by a word cloud and labeled by Llama 3 reading five randomly sampled representative sentences per cluster. Sentence-level topic modeling was conducted hierarchically in two stages. In the first stage, sentences were clustered using HDBSCAN to identify primary topic clusters. In the second stage, super-clusters were derived by applying a second round of HDBSCAN to the primary cluster centroids, grouping related primary topics into broader thematic categories. Super-cluster labels were generated by Llama 3 reading the set of primary cluster labels contained within each super-cluster. Both levels of the hierarchy are displayed in the legend labeled Scene Hierarchy accompanying each figure, with primary clusters shown as color shades of their parent super-cluster color.

Topic similarity at the report level was assessed using a best-match approach, building from prior work (Adam & Kogler, 2024; Ermakova et al., 2025; Sanz et al., 2018). Here, for each category pair, each topic centroid (defined as the L2-normalized mean of all member report embeddings) was matched to its most similar counterpart by cosine similarity. The average best-match similarity across topics was computed for each category pair. Differences in this average across pairs were assessed with a Kruskal-Wallis test, with planned Dunn post-hoc comparisons (Bonferroni corrected) testing whether certain condition pairs were more similar than others.

### Phenomenological Trajectory Analysis

Embeddings, at both the report and sentence levels, were compared by cosine similarity to six phenomenological theme centroids derived from seed sentences in Table 1 of Dikovskaya et al. (2025) (Dikovskaya et al., 2025): Physical Sensations (e.g., “body started vibrating”), Visual Alterations (e.g., “visual disturbances appeared”), Time Dilation (e.g., “feels like an eternity”), Self Dissolution (e.g., “as if I were dissolving”), Metaphysical Experience (e.g., “inner dimension at the core of matter, time, space, and thought”), and Personal Growth (e.g., “to maintain that feeling of self-compassion and self-care”). Theme centroids were computed as the mean embedding of the seed sentences associated with each theme. Mean theme scores per category were compared using Kruskal-Wallis tests with Dunn post-hoc comparisons (Bonferroni corrected). Spearman rank correlations assessed replication of the predicted phenomenological trajectory across categories. The Baseline sample was included in all trajectory analyses as a reference condition. This analysis was conducted at the report level only, as the trajectory framework was developed and validated at the level of full experiential narratives.

**Table 1.**
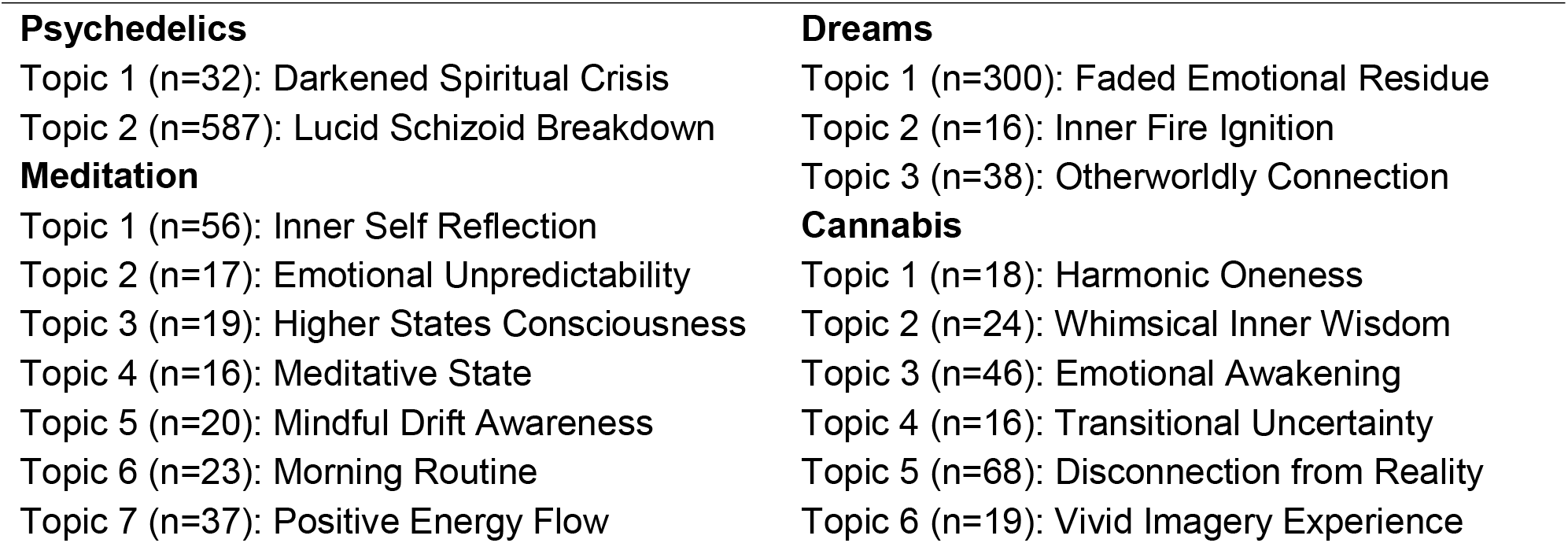
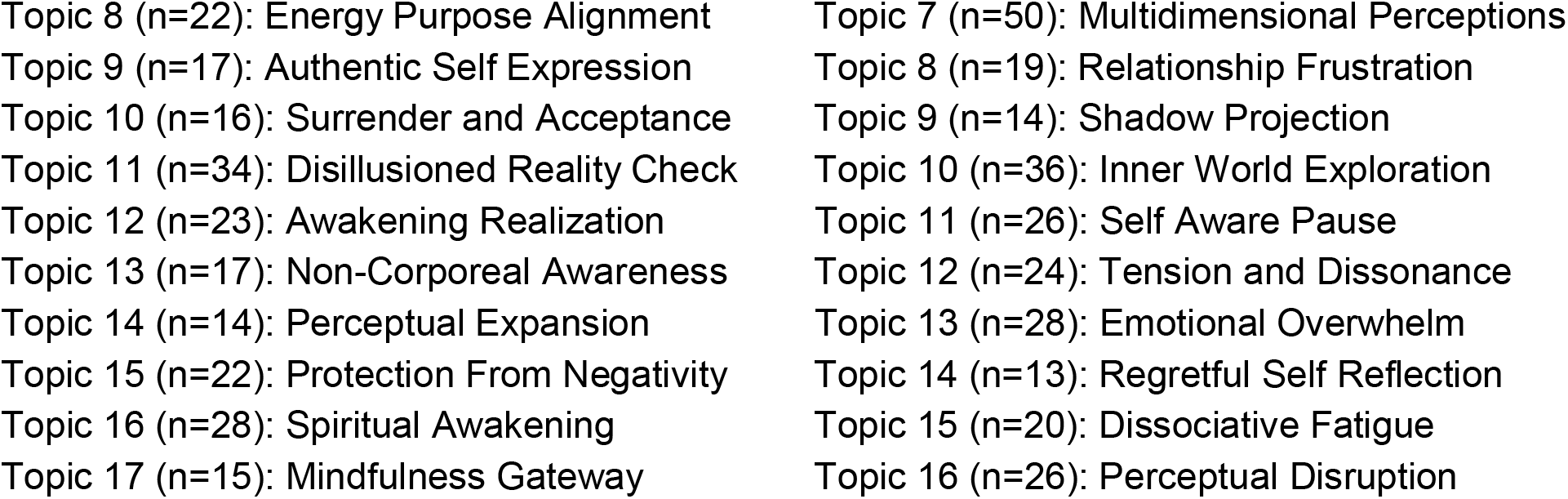
Report-level topic modeling per category. HDBSCAN-derived and Llama 3-labeled.

### Software

All analyses were conducted in Python 3.11.14 using pandas, numpy, sentence-transformers, umap-learn, hdbscan, scikit-learn, scipy, scikit-posthocs, plotly, matplotlib, and wordcloud. Random seeds were fixed at 42 throughout. Code is available at https://github.com/CMurray90/ALTERD.

## Results

### Consciousness state space

Across the ALTERD categories of Psychedelics, Cannabis, Meditation, and Dreams, a three-dimensional UMAP projection of all 2,242 reports revealed a spatial organization comprised of distinct clusters associated with each category (**Fig. 1**). To address the emergent framework of this spatial organization, we used a language model (Llama 3) to label the three axes of the space. This resulted in the labels Disintegration ↔ Integration, Desiccation ↔ Revitalization, and Desolation ↔ Abundance. To gain more insight into these labels, we used Llama 3 to generate an additional set of labels made of 2-3 words for each axial pole (**Supplementary Table 1**). The Psychedelics cluster was positioned near to the Disintegration edge, alongside Dreams, whereas the Cannabis cluster was positioned near to the Integration edge, alongside Meditation. A fifth cluster, containing a subset of reports across all four categories, was positioned between Disintegration and Integration, near to the edges of Revitalization and Abundance.

**Figure 1.**
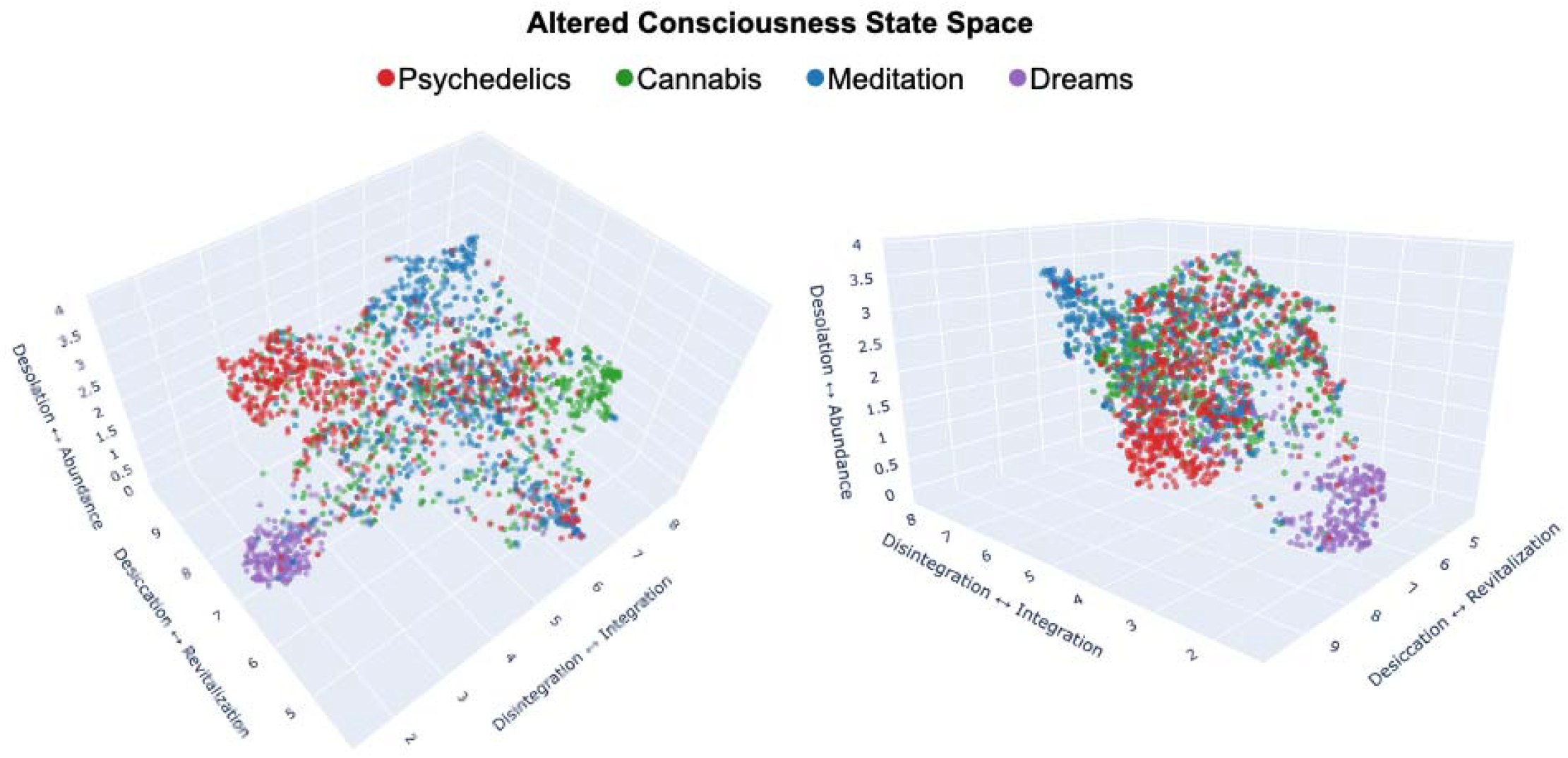
Three-dimensional UMAP projection and “state space” of ALTERD reports. Each point represents one first-person report (Psychedelics n=620, Cannabis n=620, Meditation n=641, Dreams n=361), colored by category. Two viewing angles are shown. Axes reflect emergent phenomenological dimensions of the consciousness state space, labeled by Llama 3 from 8 reports at each dimensional extreme, presented as [low value] ↔ [high value]: Disintegration ↔ Integration, Desiccation ↔ Revitalization, and Desolation ↔ Abundance. UMAP random_state = 42.

Word clouds indicating the relative frequency of words included in the reports from each category are shown in supplementary material (**Supp. Fig. 1**).

### Semantic similarity

Across the four categories, cosine similarity revealed the degree of semantic overlap in reports from each category (**Fig. 2A**). Specifically, the Dreams-Psychedelics pair was the most similar pair of categories, while Dreams-Cannabis was the least similar pair. The Dreams category was also the most self-similar category, while the Cannabis category was the least self-similar.

**Figure 2.**
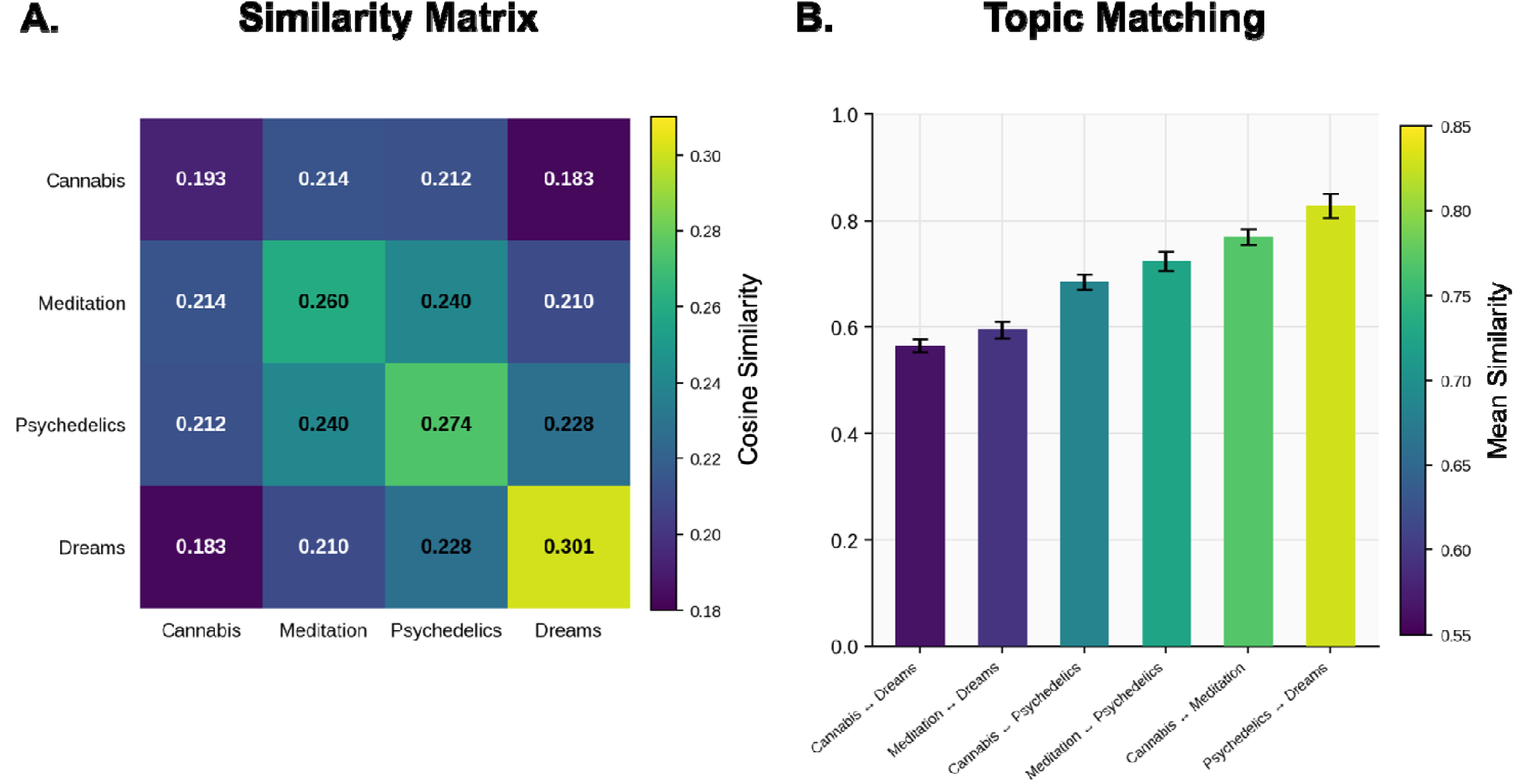
Semantic similarity across altered states of consciousness. Panel A shows mean pairwise cosine similarity between report embeddings across altered states categories. Red color indicates most similar pairs and green indicates least. Diagonal cells reflect internal cohesion (within-category similarity); off-diagonal cells reflect between-category similarity (Cannabis n=620, Meditation n=641, Psychedelics n=620, Dreams n=361; total N=2,242). Panel B shows mean best-match topic similarity across report categories, where HDBSCAN-derived topic centroids from one category were matched to their closest counterpart by cosine similarity. Topic counts were: Cannabis=16, Meditation=17, Psychedelics=2, Dreams=3 (see Table 1 for labeled topics). Bars indicating best-match topic similarity are sorted ascending and error bars represent ±1 SEM. Kruskal-Wallis test confirmed significant differences across pairs (H=27.017, p<0.001); Dunn post-hoc comparisons between pairs (Bonferroni corrected) are reported in text.

We then conducted topic modeling for each category, and the topics were subsequently labeled by Llama 3 (**Table 1**). Pairs of categories were then analyzed at the topic level through a best-match similarity analysis of topic centroids (**Fig. 2B**). We found that across category pairs there was a significant difference in the degree of similarity (Kruskal-Wallis: H=27.017, p<0.001). Post-hoc pairwise comparisons further revealed significant differences between pairs, indicating that the topics between Meditation and Cannabis were significantly more similar than topics between Dreams and Cannabis or Meditation and Dreams (ps<0.001). Meaningful statistical comparisons related to the Psychedelics topics were precluded by the low number of topics derived from HDBSCAN (2 topics) in this category.

To control for individual differences, we additionally examined semantic similarity in a within-subjects subset of ALTERD users who contributed at least one report to each of the four categories. For each user, one report per category was randomly selected for analysis (n=21 users, 84 reports total; **Supp. Fig. 2A**). This within-subjects approach reproduced findings reported above, including that the Dreams-Psychedelics was the most similar pair, that the Dreams category was the most self-similar, and that the Cannabis category was the least self-similar. We then applied the same topic-level best-match analysis to the 84 reports to examine differences in topic similarity between category pairs, but found no significant differences across categories (**Supp. Fig.2B**).

### Visual phenomenology

As a next step, we sought to address the nature of fine-grained phenomenological and perceptual content across the categories using the full set of reports from each category. We therefore moved from report-level embeddings to performing a sentence-level analysis of extracted visual scene sentences. The sentence embeddings were visualized in three-dimensional space (UMAP) for each category and hierarchical topic modeling identified topic clusters with distinct perceptual themes (**Fig. 3**).

**Figure 3.**
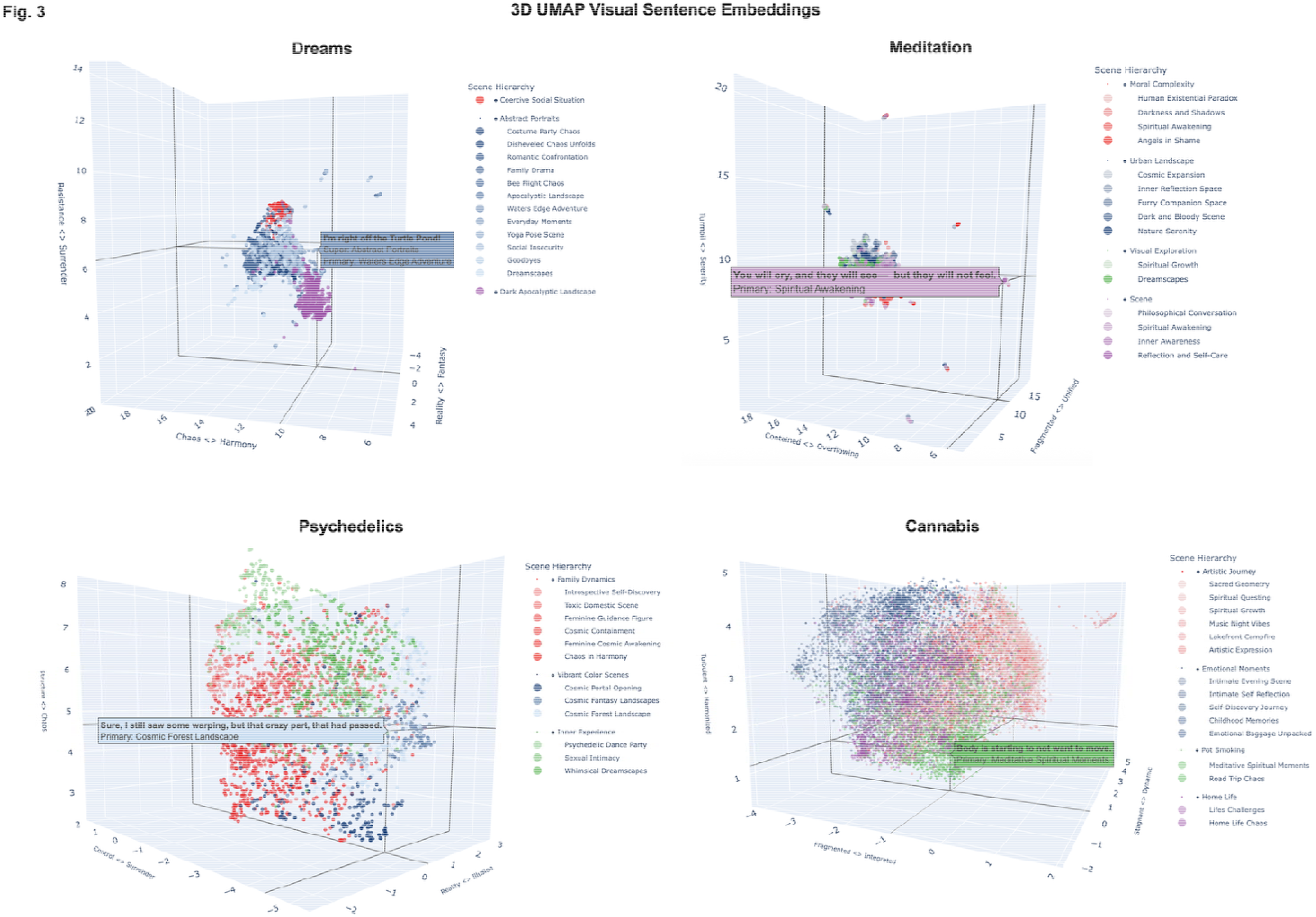
Three-dimensional UMAP projections of visual scene sentences across altered state categories. Each panel shows a 3D UMAP of visual scene sentences extracted from reports in that category along with a representative sentence. Points reflect individual sentences, colored by topic cluster. Super-clusters (top-level groupings) and their constituent primary clusters are shown in the Scene Hierarchy legend to the right of each UMAP projection. Topic labels were generated by Llama 3 reading representative sentences from each cluster presented as [low value] ↔ [high value]. UMAP random_state=42.

To quantify the similarities between categories at this sentence level, we performed two analyses. First, cosine similarity indicated that globally, the visual scene sentences in Meditation and Cannabis were the most similar pairing, whereas Meditation and Dreams were the least similar (**Fig. 4A**). Second, a nearest-neighbor overlap analysis further revealed this same pattern (**Fig. 4B**), together indicating that at both a global and local structural level, Meditation and Cannabis share the most overlap in visual phenomenology amongst the four category pairs.

**Figure 4.**
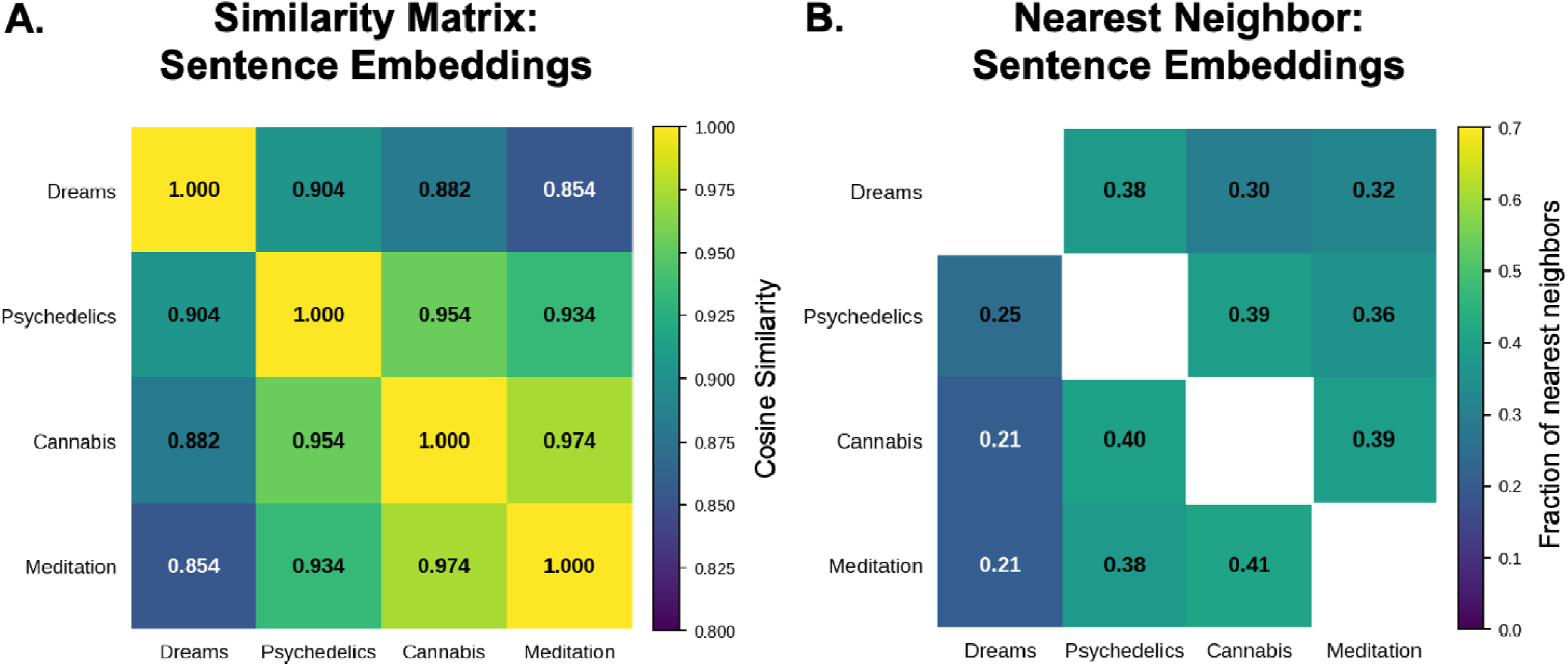
Visual scene sentence similarity across altered state categories. (A) Cosine similarity matrix showing similarity between the mean visual scene sentence embeddings for each category at the global level. Each cell reflects the similarity between two centroids computed from all visual scene sentences in that category, with embeddings L2-normalized prior to computation. (B) Cross-category nearest neighbor overlap matrix showing the fraction of each category’s visual scene sentences whose nearest neighbor belongs to another category. Each cell reflects the proportion of sentences from one category whose nearest neighbor belongs to another category.

### Altered trajectory

At both the report and sentence levels, we addressed how ALTERD categories mapped onto ASC experiences from baseline experience using *a priori* themes from a recent thematic analysis of ASC (Dikovskaya et al., 2025). Here, we included a fifth ALTERD category, Baseline, as an additional comparator. We found significant differences between the ALTERD categories (Baseline, Psychedelics, Cannabis, Meditation, Dreams) with respect to their degree of similarity to each theme derived from our prior thematic analysis (Physical Sensations, Visual Alterations, Time Dilation, Self Dissolution, Metaphysical Experience, and Personal Growth) (Kruskal-Wallis, all themes p<0.001; **Fig. 5A, Fig. 5B**). Baseline reports were the least similar to the altered states themes overall, whereas Psychedelics and Meditation reports were the most similar to altered states themes overall. Post-hoc Dunn comparisons confirmed significant pairwise differences between ALTERD categories with respect to the *a priori* altered states themes (**Supplemental Table 2**).

**Figure 5.**
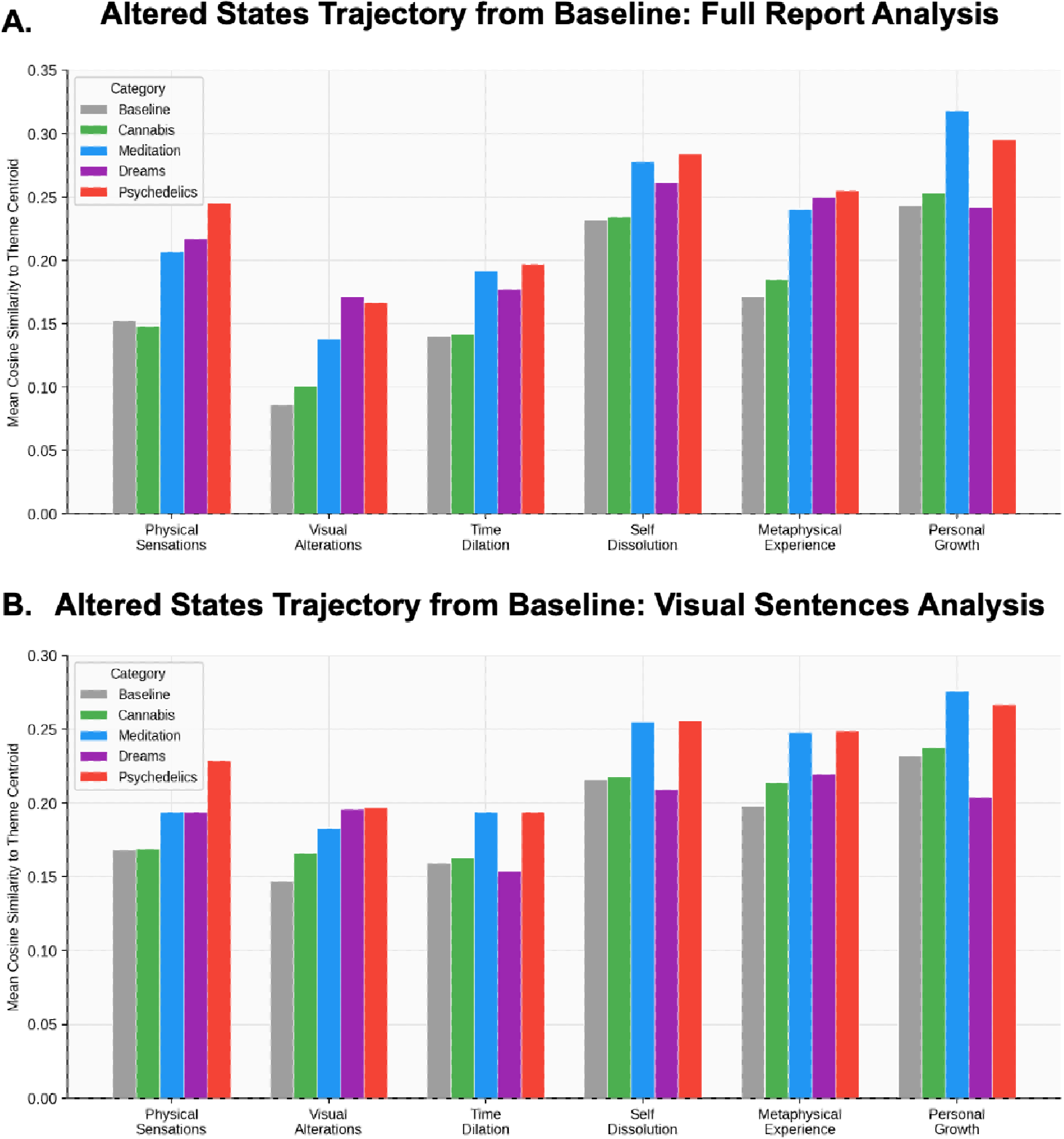
Mean cosine similarity in (A) full report embeddings and (B) visual sentence embeddings to phenomenological theme centroids across altered state categories. Each bar shows the mean cosine similarity between embeddings in a given category and the centroid of each phenomenological theme, derived from seed sentences in Table 1 of Dikovskaya et al. (2025). Themes represent a trajectory of deepening altered state experience: Physical Sensations, Visual Alterations, Time Dilation, Self Dissolution, Metaphysical Experience, and Personal Growth. Baseline (gray) reflects reports from ALTERD users prior to altered state experiences (n=18,179). All themes differed significantly across categories (Kruskal-Wallis, all p<0.001).

## Discussion

Here, we analyzed data from ALTERD, a large-scale mobile application that captures reports of ASC including Dreams, Meditation, Cannabis, and Psychedelics. We examined similarities and differences in natural language between these categories, both at the level of the full report and at the level of individual sentences related to visual phenomenology. We also compared these categories and the ALTERD Baseline category to a recently published trajectory of ASC from baseline (Dikovskaya et al., 2025), which identified six emergent themes related to the trajectory of altered states. Our aim in the current manuscript was to test the hypothesis that the phenomenology of serotonergic hallucinogens, also known as psychedelics (Nichols, 2016), includes an affordance of insight (i.e., are “mind-manifesting”) along with their perception-distorting nature. While, as we discuss below, we recognize these are broad categories, in the present work we operationalize the experience of genuine inner observation and insight as similarity to the experience of meditation and we operationalize the experience of illusory inner-generated perception as similarity to the experience of dreaming. In addition, the inclusion of Cannabis served to address its phenomenological character relative to Psychedelics for greater context and rigor.

Overall, we find the phenomenology of Psychedelic experiences to be most similar to that of Dreams, as assessed by cosine similarity of full reports and topic modeling, while also exhibiting, alongside Meditation, the highest similarity to the Personal Growth theme. Conversely, we find the phenomenology of Cannabis experiences to be most similar to that of Meditation, as assessed by cosine similarity of full reports and topic modeling, while exhibiting, alongside Baseline, the lowest similarity to the Personal Growth theme.

With respect to the global structure of the consciousness state space, we find the four experience categories to be clearly separated in a three-dimensional space, suggesting that the categories represent phenomenologically distinct, yet relationally structured, phenomenological content. As assessed with a Large Language Model (LLM), the consciousness state-space implicit in the data appears to be organized along dimensions of experiential coherence (from integration to disintegration), vitality (from revitalization to desiccation), and valence (from abundance to desolation), consistent with the themes identified in previous ASC work (Bayne, 2004; Dittrich, 1998; Revonsuo et al., 2009). The Disintegration ↔ Integration axis closely mimics a major theme in the neuroscience of consciousness. Neuroimaging studies have identified greater network integration following psychedelics associated with psychological insight scores (Carhart-Harris, 2026) and recent work has further identified an Integration-Segregation axis of neural networks across decreasing states of consciousness including segregation-related loss of consciousness under anesthesia (Dai et al., 2026; Monti et al., 2013). We found that the Meditation and Cannabis categories sat along the Integration edge of the space, whereas Dreams sat at the Disintegration edge.

We then examined semantic and topic similarity at the level of full reports and at the level of individual sentences related to visual phenomenology. Across both the full report and sentence levels, the closest pairing was Psychedelics-Dreams, suggesting that psychedelic experiences share the idiosyncratic and bizarre quality of dreaming rather than with the mindful, inwardly directed quality of meditation. This interpretation is consistent with prior work (Sanz et al., 2018) and supported by topic modeling at both the full report and sentence levels, with Psychedelics yielding topic labels including “Fantasy Landscapes” and “Whimsical Dreamscapes,” similar to the “Otherworldly,” “Abstract Portraits,” and “Apocalyptic Landscapes” from Dreams. Cannabis and Meditation reports were significantly more similar to one another than either was to Dreams, suggesting that Cannabis and Meditation may share a common phenomenological substrate, one oriented toward a salient present-moment awareness and internal reflection, which in turn may produce intersubjective observations, as opposed to idiosyncratic hallucinations. This is interpretation is consistent with prior work (Murray & Srinivasa-Desikan, 2022) and supported by topic modeling at both the full report and sentence levels, with Cannabis yielding labels including “Harmonic Oneness,” “Inner Wisdom,” “Inner World Exploration,” “Spiritual Questing,” and “Spiritual Growth,” similar to the “Inner Self Reflection,” “Awakening Realization,” “Inner Awareness,” “Spiritual Awakening” and “Spiritual Growth” from Meditation. To further illustrate this point, representative quotes from the Cannabis topic labeled as “Harmonic Oneness” read:

> *“This… is the fundamental fabric of reality, weaving together space, time, and consciousness into a unified, interconnected whole*.*”*

> *“*Φ *(Phi): The golden ratio, representing the intrinsic harmony and balance that underlies all of existence*.*”*

Despite pairings between Psychedelics-Dreams and Cannabis-Meditation, when we examined *a priori* themes of ASC trajectory from baseline, we found that, Psychedelics, and not Cannabis, tightly coupled with Meditation, particularly with respect to Personal Growth. This finding underscores the potential therapeutic uses of psychedelic medicines, and not cannabis, for transformative and enduring psychiatric benefits (Jylkkä et al., 2025; Vargas et al., 2021; Yaden & Griffiths, 2021). The apparent juxtaposition between dream-like phenomenology and personal growth after psychedelics lends nuanced support for the psychedelic hypothesis—that serotonergic hallucinogens have a mind-manifesting character linked to therapeutic outcomes—despite also inducing an acute, hallucinogenic, dream-like character. To illustrate further, representative quotes from the Psychedelics category associated with Personal Growth read:

> *“This was an incredible healing and mystical experience*.*”*
>
> *“I was able to forgive everyone, I felt guilty for things without feeling bad about it. I was able to forgive myself. I was able to see the world through someone else’s eyes, feeling empathy and a desire to forgive*.*”*

The current findings using ALTERD reports represents the latest step in a growing line of work investigating ASC using online sources, including Erowid and Reddit (Baggott et al., 2010; Coyle et al., 2012; Dikovskaya et al., 2025; Hase et al., 2022; Lawrence et al., 2022; Martial et al., 2019; Zamberlan et al., 2018). Earlier analyses of Erowid and Reddit reports identified frequently reported words such as “reality,” “dimension,” “universe,” and “consciousness” as hallmarks of ASC (Coyle et al., 2012), while others distinguished discrete phenomenological profiles across substances, linking mystical experiences to DMT and emotional or cognitive language to MDMA (Hase et al., 2022). In comparing dreams to 165 drug-induced states, the psychedelic LSD and deliriant datura were found to be most similar to dreaming (Sanz et al., 2018), whereas of the same 165 substances, the dissociatives salvia and ketamine were found to be most similar to near-death experiences (Martial et al., 2019). In a recent qualitative analysis, salvia and ketamine reports were also found to indicate a sense of authentic observations related to mind or being, while portraying an emergent phenomenological structure (Dikovskaya et al., 2025). From ALTERD, full text reports labeled as “salvia” (n=6 non-fragment reports) are included in Supplementary Material, which support this previously reported spindle-like mechanistic structure:

> *“I’m looking at reality from the third person. Like sticking my head out of a window. Though in this case the window of reality itself. It’s moving, twisting, rotating and leaving colourful vibrant trails behind it. I realise this is it. No mystery. No cosmic revelation. Just the bare mechanics of everything there is*.*”*

> *“It’s like I’ve slipped out of this reality and I could feel a cylindrical motion around me. I felt like I was a long 3D structure of a cylinder in which I was in the centre, the spindle of Life and this reality is a slice of this cylinder. Think of it like a pie chart with so many different thin slices. This reality is just one of those slices and there are millions more*.*”*

The present study advances prior work by replacing retrospectively sourced, publicly available forum posts with prospectively submitted reports from ALTERD, a purpose-built mobile application whose users self-document experiences across designated ASC categories including Dreams, Meditation, Cannabis, and Psychedelics. This shift to a mobile application-based reporting platform addresses key limitations, including recall bias arising from the temporal distance between experience and documentation, narrative shaping introduced by writing for a public audience, and the absence of standardized metadata on substance category and contextual variables. The ALTERD dataset, with thousands of reports, provides the scale necessary to conduct within-subjects analyses, hierarchical sentence-level topic modeling, and three-dimensional consciousness state space mapping that were not feasible in prior Erowid- and Reddit-based investigations, while preserving the ecological validity that has made online report corpora valuable to the phenomenology of consciousness literature.

Several limitations of the present study warrant consideration. First, our investigation rests on two key pillars, which is the use of meditation and dreaming as opposite comparators for addressing authentic and illusory perceptions, respectively, during altered states of consciousness. However, this framework has not been established previously. Moreover, the ALTERD Meditation category contains various types of meditations, most of which are not specified, and it is unclear which form of meditation, if any, is most reliable as a positive control for generating authentic insights and observations of mind. Notably however, we found that the Meditation and Dreams categories were often orthogonal in our analyses, helping to validate the use of these categories to test the validity of the mind-manifesting, psychedelic nomenclature. Second, the data analyzed in this study were obtained naturalistically though a mobile application, introducing a number of potential confounds, from self-selection bias to the role of self-assigned category labeling and self-monitored responses. The data were analyzed without verification of either the category or prompt to specifically report on the acute experience associated with that category. Together, these limitations suggest that findings should be interpreted as reflecting the structure of reported altered state phenomenology within a self-selected population. Future work should seek to complement large-scale naturalistic data with controlled, prospective designs that permit verification of substance administration, standardization of assessment timing, and sampling across more diverse populations.

Here, we used ALTERD reports from Psychedelics, Cannabis, Meditation, and Dreams to test the construct validity of the psychedelic hypothesis that serotonergic hallucinogens are mind-manifesting. Across multiple analyses, psychedelic experiences resembled more closely Dreams than Meditation in their phenomenological profile, suggesting that their perceptual character is predominantly dream-like/hallucinogenic. At the same time, psychedelics aligned with Meditation in the thematic trajectory of ASC, particularly with respect to Personal Growth. These findings suggest that serotonergic hallucinogens possess a hallucinogenic profile alongside some mind-manifesting characteristics related to therapeutic potential. This interpretation is consistent with the emergence of “psychedelic therapies” and theoretical frameworks that treat serotonergic hallucinogens as amplifiers of mental content rather than generators of fixed hallucinatory states (Carhart-Harris & Friston, 2019; Letheby & Gerrans, 2017). Cannabis, meanwhile, although not aligned with Personal Growth, nonetheless emerged as phenomenologically closer to Meditation than to Dreams across multiple analyses, a finding that may reflect the use of cannabis in contemplative contexts and as a possible tool to investigate meditative states and associated internal reflections and observations. Together, our analyses support Huxley and Osmond’s psychedelic etymology for both cannabis and the serotonergic hallucinogens, but in largely contrasting ways.

---

Supplemental Table 1 shows pairwise differences from Dunn post-hoc comparisons (Bonferroni corrected).

