## Supplemental Materials for "Mind-manifesting? Examining naturalistic reports from psychedelics, cannabis, meditation, and dreams from the ALTERD mobile application"

**Supplemental Methods**

To evaluate whether reports excluded by the 100-word minimum represented “unsuitable” reports, including low-information fragments, rather than meaningful but brief reports, we developed a suitability filter in Python. A report was classified as "suitable" if it (a) was not a structural artifact (markdown formatting, survey-format content, or degenerate repeated text) and (b) contained minimal first-person experiential language (e.g., "I felt...", "I saw..."). This classifier was applied to all reports (after removing duplicates) across the four categories prior to word-count filtering (N=22,614), and separately to the final 620-report Cannabis subsample, its 4,006-report source pool, and the other three category samples, to assess whether downsampling preserved comparable report quality.

**Supplemental Results**

Of the 22,614 reports, 52.2% were classified as suitable. Reports meeting the 100-word minimum were substantially more likely to be classified as suitable (72.7%, 4,092/5,628) than those excluded by the threshold (45.4%, 7,711/16,986), χ²(1) = 1262.66, p < .001, indicating the word-count filter is associated with report quality as intended, while also acknowledging that a meaningful minority of brief reports below threshold were independently classified as substantive. Nonetheless, a printed list of example reports revealed little meaning in those that passed for “suitable.”

[Cannabis, 3 words] I feel things

[Cannabis, 2 words] I’m high

[Psychedelics, 3 words] I love tripping

[Cannabis, 3 words] I love light

[Psychedelics, 3 words] I love you

For the Cannabis downsampling, the final 620-report analytic subsample showed a "suitable" rate (72.9%) comparable to the full eligible pool it was drawn from (70.8%, n=4,006) and to the other three category samples (Meditation 80.0%, Psychedelics 77.4%, Dreams 72.6%), indicating the random downsample did not introduce a quality bias relative to its source pool or the other categories.

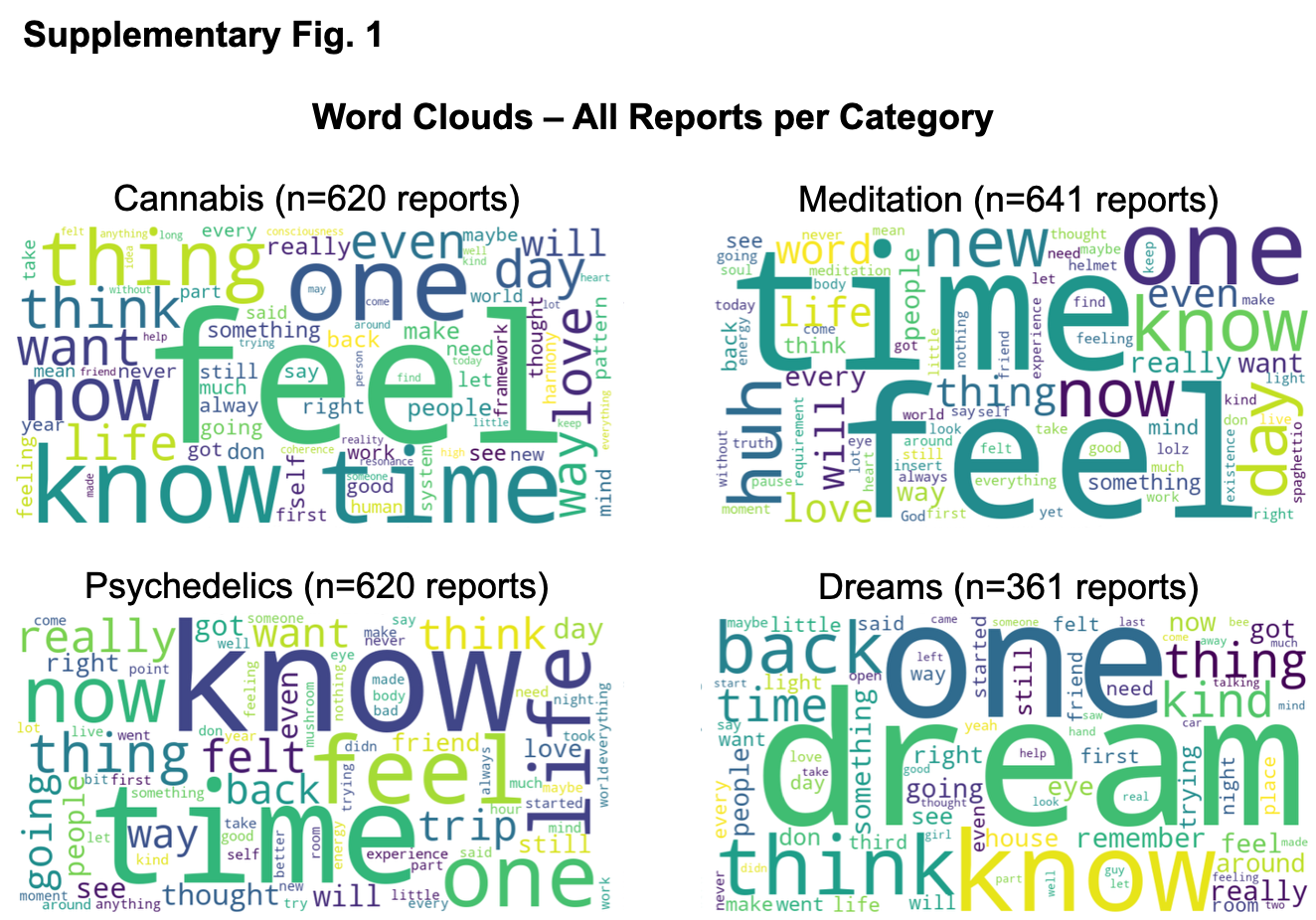

**Supplemental Figure 1.** Word clouds of reported content associated with altered states of consciousness. Each panel shows a word cloud generated from all reports within that category (Cannabis n=620, Meditation n=641, Psychedelics n=620, Dreams n=361). Word size is proportional to frequency. Common English stopwords and contraction fragments were removed prior to generation. Word clouds reflect the most distinctive lexical content of each category's reports rather than topic-level structure.

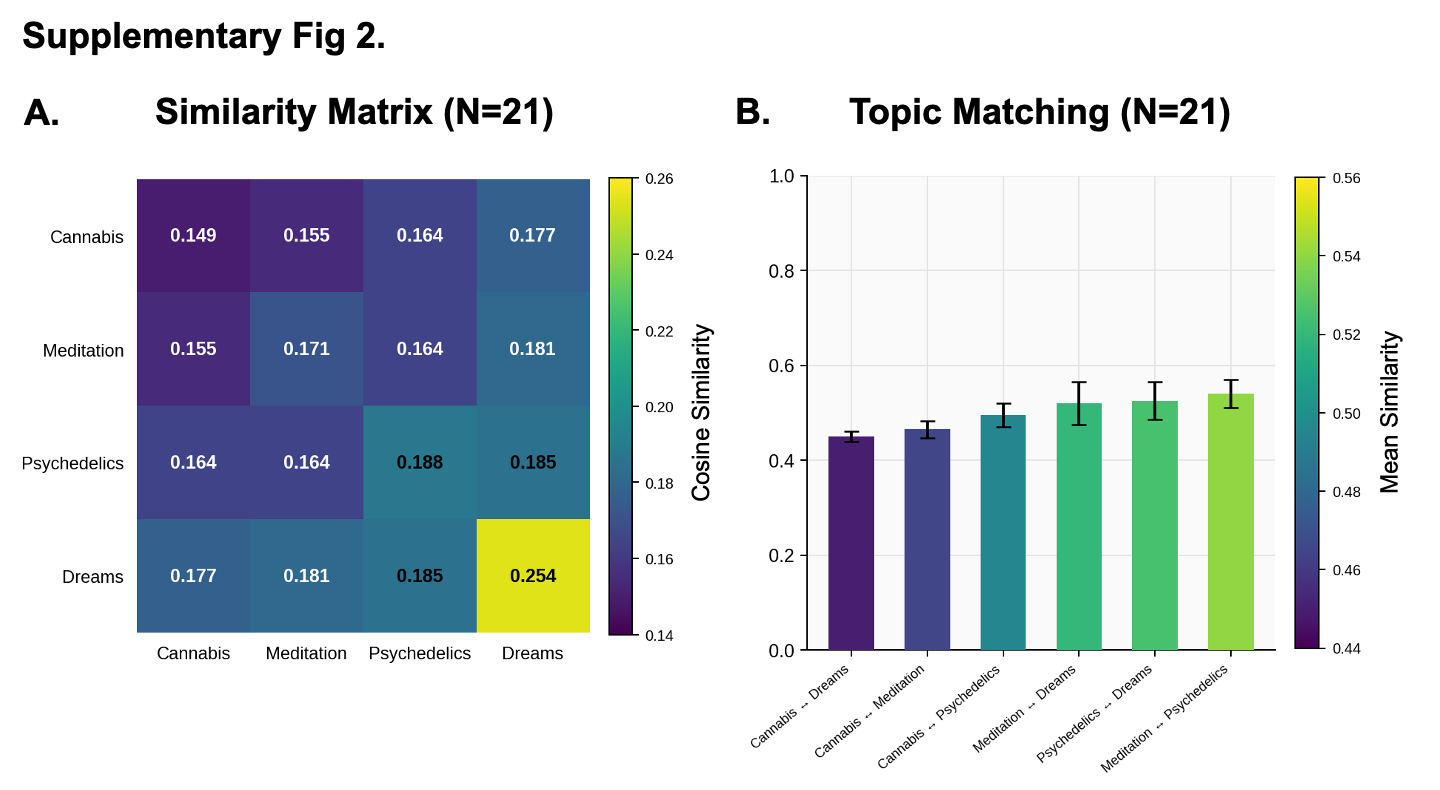

| **Supplemental Table 1** | | |  |
| --- | --- | --- | --- |
| **Axis** | **Pole** | **Label #** | **Label** |
| **Disintegration** | | | |
| Axis 1 | Low pole | 1 | Vivid Disorientation |
| Axis 1 | Low pole | 2 | Fragmented Reality |
| Axis 1 | Low pole | 3 | Disrupted Familiarity |
| Axis 1 | Low pole | 4 | Turbulent Transition |
| Axis 1 | Low pole | 5 | Unsettled Experience |
| **Integration** | | | |
| Axis 1 | High pole | 1 | Celestial Guidance |
| Axis 1 | High pole | 2 | Cosmic Insights |
| Axis 1 | High pole | 3 | Stellar Awareness |
| Axis 1 | High pole | 4 | Astrological Harmony |
| Axis 1 | High pole | 5 | Galactic Clarity |
| **Desolation** | | | |
| Axis 2 | Low pole | 1 | Essence of Absence |
| Axis 2 | Low pole | 2 | Voidal Horizon |
| Axis 2 | Low pole | 3 | Celestial Weight |
| Axis 2 | Low pole | 4 | Obsidian Silence |
| Axis 2 | Low pole | 5 | Fractured Illumination |
| **Abundance** | | | |
| Axis 2 | High pole | 1 | Relational Turmoil |
| Axis 2 | High pole | 2 | Emotional Vortex |
| Axis 2 | High pole | 3 | Unrequited Love |
| Axis 2 | High pole | 4 | Power Dynamics Tension |
| Axis 2 | High pole | 5 | Relationship Reckoning |
| **Desiccation** | | | |
| Axis 3 | Low pole | 1 | Effortless Connection |
| Axis 3 | Low pole | 2 | Reflective Relaxation |
| Axis 3 | Low pole | 3 | Mind-Expanding Mellow |
| Axis 3 | Low pole | 4 | Comforting Catharsis |
| Axis 3 | Low pole | 5 | Euphoric Exploration |
| **Revitalization** | | | |
| Axis 3 | High pole | 1 | Abyssal Fervor |
| Axis 3 | High pole | 2 | Dark Radiance |
| Axis 3 | High pole | 3 | Eternal Ignition |
| Axis 3 | High pole | 4 | Visceral Blaze |
| Axis 3 | High pole | 5 | Sacred Ferocity |

| **Supplemental Table 2.** Altered trajectory: Pairwise differences (Bonferroni corrected) | | |
| --- | --- | --- |
| Comparison | Report-level | Sentence-level |
| ***Physical Sensations*** | | |
| Baseline vs. Cannabis | p = 1.0000 (ns) | p = 1.0000 (ns) |
| Baseline vs. Meditation | p < 0.001*** | p < 0.001*** |
| Baseline vs. Dreams | p < 0.001*** | p < 0.001*** |
| Baseline vs. Psychedelics | p < 0.001*** | p < 0.001*** |
| Cannabis vs. Meditation | p < 0.001*** | p < 0.001*** |
| Cannabis vs. Dreams | p < 0.001*** | p < 0.001*** |
| Cannabis vs. Psychedelics | p < 0.001*** | p < 0.001*** |
| Meditation vs. Dreams | p = 0.1222 (ns) | p = 0.3162 (ns) |
| Meditation vs. Psychedelics | p < 0.001*** | p < 0.001*** |
| Dreams vs. Psychedelics | p = 1.0000 (ns) | p < 0.001*** |
| ***Visual Alterations*** | | |
| Baseline vs. Cannabis | p = 0.0013** | p < 0.001*** |
| Baseline vs. Meditation | p < 0.001*** | p < 0.001*** |
| Baseline vs. Dreams | p < 0.001*** | p < 0.001*** |
| Baseline vs. Psychedelics | p < 0.001*** | p < 0.001*** |
| Cannabis vs. Meditation | p < 0.001*** | p < 0.001*** |
| Cannabis vs. Dreams | p < 0.001*** | p < 0.001*** |
| Cannabis vs. Psychedelics | p < 0.001*** | p < 0.001*** |
| Meditation vs. Dreams | p < 0.001*** | p < 0.001*** |
| Meditation vs. Psychedelics | p < 0.001*** | p = 0.3877 (ns) |
| Dreams vs. Psychedelics | p = 0.9859 (ns) | p = 0.3864 (ns) |
| ***Time Dilation*** | | |
| Baseline vs. Cannabis | p = 1.0000 (ns) | p < 0.001*** |
| Baseline vs. Meditation | p < 0.001*** | p < 0.001*** |
| Baseline vs. Dreams | p < 0.001*** | p = 1.0000 (ns) |
| Baseline vs. Psychedelics | p < 0.001*** | p < 0.001*** |
| Cannabis vs. Meditation | p < 0.001*** | p < 0.001*** |
| Cannabis vs. Dreams | p < 0.001*** | p < 0.001*** |
| Cannabis vs. Psychedelics | p < 0.001*** | p < 0.001*** |
| Meditation vs. Dreams | p = 1.0000 (ns) | p < 0.001*** |
| Meditation vs. Psychedelics | p = 1.0000 (ns) | p = 1.0000 (ns) |
| Dreams vs. Psychedelics | p = 0.4243 (ns) | p < 0.001*** |
| ***Self Dissolution*** | | |
| Baseline vs. Cannabis | p = 1.0000 (ns) | p = 1.0000 (ns) |
| Baseline vs. Meditation | p < 0.001*** | p < 0.001*** |
| Baseline vs. Dreams | p < 0.001*** | p = 0.0176* |
| Baseline vs. Psychedelics | p < 0.001*** | p < 0.001*** |
| Cannabis vs. Meditation | p < 0.001*** | p < 0.001*** |
| Cannabis vs. Dreams | p < 0.001*** | p = 0.0013** |
| Cannabis vs. Psychedelics | p < 0.001*** | p < 0.001*** |
| Meditation vs. Dreams | p = 0.2748 (ns) | p < 0.001*** |
| Meditation vs. Psychedelics | p = 1.0000 (ns) | p = 1.0000 (ns) |
| Dreams vs. Psychedelics | p = 0.0659 (ns) | p < 0.001*** |
| ***Metaphysical Experience*** | | |
| Baseline vs. Cannabis | p = 0.0544 (ns) | p < 0.001*** |
| Baseline vs. Meditation | p < 0.001*** | p < 0.001*** |
| Baseline vs. Dreams | p < 0.001*** | p < 0.001*** |
| Baseline vs. Psychedelics | p < 0.001*** | p < 0.001*** |
| Cannabis vs. Meditation | p < 0.001*** | p < 0.001*** |
| Cannabis vs. Dreams | p < 0.001*** | p < 0.001*** |
| Cannabis vs. Psychedelics | p < 0.001*** | p < 0.001*** |
| Meditation vs. Dreams | p = 0.1264 (ns) | p < 0.001*** |
| Meditation vs. Psychedelics | p = 1.0000 (ns) | p = 1.0000 (ns) |
| Dreams vs. Psychedelics | p = 1.0000 (ns) | p < 0.001*** |
| ***Personal Growth*** | | |
| Baseline vs. Cannabis | p = 0.3030 (ns) | p = 0.1692 (ns) |
| Baseline vs. Meditation | p < 0.001*** | p < 0.001*** |
| Baseline vs. Dreams | p = 1.0000 (ns) | p < 0.001*** |
| Baseline vs. Psychedelics | p < 0.001*** | p < 0.001*** |
| Cannabis vs. Meditation | p < 0.001*** | p < 0.001*** |
| Cannabis vs. Dreams | p = 1.0000 (ns) | p < 0.001*** |
| Cannabis vs. Psychedelics | p < 0.001*** | p < 0.001*** |
| Meditation vs. Dreams | p < 0.001*** | p < 0.001*** |
| Meditation vs. Psychedelics | p = 0.0076** | p = 0.0025** |
| Dreams vs. Psychedelics | p < 0.001*** | p < 0.001*** |

### **ALTERD Salvia Reports — Full Text**

#### **Report 1**

Subject ID: 61413e6b-b415-537a-5537-4ca9c7e8d292	|	Timestamp: 2025-12-03T14:38:32.466000+00:00

Had my Ego-Death today. Not that bad as they say if you know how to go with it. It show‘s you the Truth because you don‘t get blinded by your emotions anymore. let‘s you filter/sort your emotions and thoughts. I can‘t even really describe this higher form of self i‘ve discovered in me. It felt acient and thoughts we‘re storming me like crazy. It opens doors to spirituality, understanding, knowledge, history etc.. It‘s a good way to open your mind. If just some people we‘re ready to let this kind of „knowledge“ enter them. Because, first i didn‘t even really wanted to do Salvia because People we‘re sayin „you‘ll live 300years as an door or something“. But my bro enlightend me what it REALLY is. It‘s an Key, an expansion on your mind. A tool to control your mind/emotions. It opened Acient doors to me, i understood the history of humans, how we work, and how we should really live. It feels like i‘m reborn.

#### **Report 3**

Subject ID: c72fa72d-9fd6-8c37-0a9e-67e70ca00b2d	|	Timestamp: 2025-08-09T18:14:56.811000+00:00

Salvia takes me to the same place it always does. The outside of everything. Not space. Not another dimension. But beyond the multiverse itself. I'm looking at reality from the third person. Like sticking my head out of a window. Though in this case the window of reality itself.

It's moving, twisting, rotating and leaving colourful vibrant trails behind it. I realise this is it. No mystery. No cosmic revelation. Just the bare mechanics of everything there is.

I can see the multiverse. Millions of realities just like this one and more of me doing the exact same thing.

There are also entities. All encompassing. They regard me with a mild and mocking amusement. Knowing I can never explain all this once I'm back.

#### **Report 4**

Subject ID: c72fa72d-9fd6-8c37-0a9e-67e70ca00b2d	|	Timestamp: 2025-07-13T20:10:15.239000+00:00

Life starts sliding away, and going around in anti clockwise circles at a steady rate. It's like I've slipped out of this reality and I could feel a cylindrical motion around me. I felt like I was a long 3D structure of a cylinder in which I was in the centre, the spindle of. Life and this reality is a slice of this cylinder. Think of it like a pie chart with so many different thin slices. This reality is just one of those slices and there are millions more. It was like when Microsoft Windows would crash, and you're dragging a window, and it smears behind leaving a trail of itself. That's what I felt was happening. With everything! Not just what I see or what I feel. I mean literally everything!

#### **Report 5**

Subject ID: c72fa72d-9fd6-8c37-0a9e-67e70ca00b2d	|	Timestamp: 2025-07-11T11:38:25.008000+00:00

You know when you're so engrossed and focussed in a game, a book, or a movie. You forget about your actual life and who you are. Well, that's what happened. Except I was engrossed in what we call being alive. This physical reality, our consciousness, our ego! It didn't feel like I was dying. I was just waking up, but I wasn't leaving this reality behind. This reality was ceasing to exist because it was just something I had imagined.

#### **Report 6**

Subject ID: 6ac51c7f-66b6-9bbf-f452-72727daf24fe	|	Timestamp: 2025-09-23T17:52:09.353000+00:00

I was at my friend's and he offered me some saliva to smoke. I tried it. It was a quick feeling. I dont know if I did much but I just sat there and felt heavy and yet not in my body. Maybe the soul of mine is heavy. I wonder how that would effect me here and after. I was in and out if my body like disconnected. Only for a few minutes. I breathed with all matter around me. Nothing crazy happened because I can naturally sit anywhere and feel the matter and particles in, on, and around me. Normally all I have to do is sit and relax and I notice that space between. Then I felt warm and hearing was a little muffled. But then it stopped i was good and life continued forward.

#### **Report 10**

Subject ID: c72fa72d-9fd6-8c37-0a9e-67e70ca00b2d	|	Timestamp: 2025-07-09T22:38:36.540000+00:00

On a Salvia Divinorum Trip

I was literally on the outside of everything. Beyond spiritual realms. Beyond belief systems. Beyond religion. Beyond the multiverse. Outside of existence as we know it.

But it wasn't a mystical spiritual place. It was basic and mundane.

A simplistic machine containing our reality and millions more in a boring room.

It felt without a doubt that this was it. This is the real deal. It's so obvious. How could I have forgotten? I know this, I've been here before many times. But a sobering realisation, that if this is life after death. Then I would rather be alive. It didn't seem like a great place.
